# Identification Of Protein Cargo in Extracellular Vesicles from Macrophages in Progressing and Regressing Tumors

**DOI:** 10.1101/2025.11.05.685429

**Authors:** Yining Zhu, Jiaxi Cai, Raymond T. Suhandynata, Jack D. Bui

## Abstract

Exosomes are extracellular vesicles (EVs) that carry bioactive molecules from a cell of origin and may alter gene expression of the acceptor cell via binding to cell surface receptors and/or delivering their cargo into the cell. Tumor-associated macrophages (TAMs) can increase or decrease tumor growth through various mechanisms, but the impact of TAM-EVs on tumors has been difficult to elucidate due to challenges in isolating and culturing TAMs for EV purification. In this study, we set out to uncover the protein identities within EVs from TAMs in progressing and regressing tumors, thereby uncovering EV proteins’ roles in TAM-tumor crosstalk. TAMs were purified from tumors using magnetic bead isolation and cultured for up to 9 days. TAMs from regressing tumors were found to be more M1-like by high expression of major histocompatibility complex (MHC) class II, while TAMs from progressing tumors are more M2-like (low expression of MHC class II). Long-term in vitro culture of TAMs resulted in reduced expression of MHC class II. EVs were harvested from plated TAMs, and EV proteins were identified by mass spectrometry and compared to EV proteins from bone marrow-derived macrophages (BMDMs). Unsupervised hierarchical clustering studies revealed that proteins from TAM-EVs converge based on their in vitro culture duration rather than their tumor origin. Signature proteins in TAM-EVs were characterized, with the identification of galectin-3-binding protein as a signature protein present in EVs from M1-like BMDMs and regressor macrophages. Finally, immunostimulatory pathways were identified in progressor-TAM-EVs through protein-protein interaction network analysis.

## Introduction

Exosomes, a type of extracellular vesicle (EV) ranging from 30 to 150 nm in diameter, mediate cell-cell communication by transporting bioactive molecules from one cell to another (1–3). Biogenesis of exosomes begins with endosomes, the earliest precursor of exosomes, which, after the formation of intraluminal vesicles that compartmentalize the loaded cargo, transform into multivesicular bodies that fuse with the plasma membrane, allowing exosomes with packaged cargo to bud off (4, 5). Exosomes contain biomolecules that may be distinct from the cytoplasmic content of the source cell. Upon fusing with the recipient cell membrane, the delivered cargo has the potential to reprogram distinct functions of the acceptor cell (6, 7).

Macrophages are innate immune cells that mature from short-lived blood monocytes when they extravasate into tissues (8, 9). Macrophages can polarize towards a pro-inflammatory M1 state via classical activation mechanisms by cytokines such as lipopolysaccharide (LPS) and interferon-γ (IFN-γ) (10, 11). By opposition, alternative polarization through the cytokines IL-4 or IL-13 drive macrophage differentiation to M2, which mediates anti-inflammatory responses (12, 13). Tumor-associated macrophages (TAMs) are the macrophages in the tumor microenvironment (TME) that have potential to shape the tumor niche (14, 15). Single-cell RNA sequencing of TAMs has revealed their heterogeneity and implicate that TAMs take on a spectrum of states rather than a dichotomous model of M1 and M2 macrophages. Indeed, TAMs can express both M1 and M2 genes and depending on the ratio between M1/M2 gene expression, can be termed M1-like or M2-like (16, 17).

Past research on EVs in the TME has focused on tumor-derived EVs (18–20) and their ability to promote tumor spread by creating a metastatic niche in the target organ (21). Studies on tumor-derived EVs and their cargo have demonstrated both their immunostimulatory (22–24) and immunosuppressive functions (25, 26). For instance, Marton et al. showed that melanoma-derived exosomes play anti-tumoral roles via induction of CD4^+^ T cell proliferation and stimulation (22–24), whereas Leary et al. reported that melanoma-derived exosomes, induced CD8^+^ T cell apoptosis and promoted tumor progression (25, 26).

Given the abundant studies of EVs derived from tumor cells and the critical role that TAMs take in both forming and shaping the TME, the possibility that TAMs might produce EVs to mediate crosstalk in the TME has gained interest (27, 28). Due to difficulties in harvesting TAMs directly from tumors and maintaining their viability in vitro for their EVs to be collected (29), previous studies have turned to harvesting EVs from TAM-mimics, macrophage cell lines co-cultured with tumor cells (30–32). Other studies characterized EVs obtained from bone marrow derived macrophages (BMDMs) and not TAMs (33, 34). In one publication, Cianciaruso et al. studied EV cargo from tumor suspensions containing macrophages and depleted of macrophages to infer TAM-EV protein cargo, demonstrating that TAM-EVs may possess immunostimulatory functions that are opposite to the phenotypes found in TAMs themselves (35).

Herein, we purified and plated TAMs directly from mouse tumors for up to 9 days and harvested their EVs to identify their protein cargo at various time-points (e.g. short-term, overnight, and long-term). We examined TAMs purified from progressing and regressing tumors (36) and compared their EV protein cargo with the protein cargo from EVs obtained from control murine M0/M1/M2 BMDMs. Based on unsupervised hierarchical analysis of their protein content, we found that TAM-EVs clustered based on the duration of culture in vitro rather than the nature of their original TMEs. We then characterized the proteins from progressor- and regressor-TAM-EVs and identified their signature proteins. We also determined signature proteins for EVs from M1 and M2 BMDMs and found one protein, galectin-3-binding protein (LGALs3bp), to be shared between the signature EV proteins in regressor-TAMs and M1 BMDMs. Finally, we anaylzed protein-protein interaction networks in EVs of progressor- and regressor-TAMs and found that progressor TAM-EVs were enriched in more pro-inflammatory and immunostimulatory pathways.

## Methods

### Animals

C57BL/6 mice (2-6 months of age) were utilized for experiments. For tumor experiments, only male mice were utilized since the 9609 cell line is derived from a male mouse and would be rejected if transplanted into female hosts. For bone marrow experiments, both male and female mice were used.

### Tumor cell lines and culture

The 9609 cell line was derived from a fibrosarcoma that developed in a male C57BL/6 mouse injected with the carcinogen 3’methylcholanthrene (37). This cell line grows progressively when transplanted into syngeneic wild-type (WT) mice and forms fibrosarcomas. We previously found that transducing 9609 parental cells with the hMEK1 gene induced a regressor growth phenotype (36). Control 9609-parent (progressor), 9609-GFP (progressor), and 9609-hMEK1 (regressor) cells were cultured in RPMI 1640 media (Gibco^TM^ 11875093) supplemented with 10% FBS (Peak Serum PS-FB4), 5% sodium pyruvate (Gibco^TM^ 11360070), 5% non-essential amino acids (Gibco^TM^ 11140050), 1% sodium bicarbonate (Gibco^TM^ 25080094), 1% penicillin streptomycin (Gibco^TM^ 15140122), 1% L-glutamine (Gibco^TM^ 25030081) and 14.3 mM beta-mercaptoethanol (Gibco^TM^ 21985023). Cells were grown in incubators supplied with 5% carbon dioxide and maintained at 37 degrees Celsius.

### Tumor injection, isolation, and dissociation

Tumor cells were detached from tissue culture flasks via trypsinization and washed three times in cold HBSS (Gibco^TM^ 14175079) with calcium and magnesium and injected at 1-5 million cells in the subcutaneous space of a syngeneic mouse (36). Mice bearing tumors with areas greater than 64 mm^2^ (21-55 days after transplant) were euthanized and tumors were harvested. Tumors were dissociated first mechanically using sterile razor blades and then enzymatically via type 1A collagenase (1 mg/mL) (Worthington Biochemical Corporation LS004197) digestion for 30 minutes at 37 degrees Celsius. The tumor single-cell suspension was passed through a 100 μm filter and spun into a tumor pellet for purification of TAMs.

### TAM purification

TAMs were purified using positive magnetic bead selection (STEMCELL Technologies). The tumor pellet was resuspended in buffer (PBS with 2% FBS and 1 mM EDTA) and incubated with primary F4/80 antibodies followed by secondary antibodies conjugated with metal beads. F4/80-positive cells were selected using a magnet and washed twice in serum-free media to remove any serum-derived exosomes. 100,000 to 200,000 cells were seeded into each well of a TC-treated 24-well plate, with serum-free exosome-free media (RoosterCollect™-EV, RoosterBio) for up to 9 days for exosome harvest.

### Fluorescence-Activated Cell Sorting (FACs)

Cells were incubated with Fc-block (2.4G2) and stained with antibodies to detect F4/80, CD11b, and MHC class II (I-E/I-Ab) (Biolegend, San Diego). Dead cells were identified by 7-actinomycin D (7-AAD) staining. We characterize TAMs to be F4/80+, CD11b-high and 7-AAD-low. MHC class II was used to assess the M1-like character in the TAMs.

### Bone-marrow derived macrophages (BMDMs)

C57BL/6 mice were euthanized, and their femur and tibia were isolated. Bone marrow was obtained through mechanical flushing with dPBS using a 22G needle. After the cells were filtered using a 100 μm filter, they were suspended in DMEM (Gibco^TM^ 11965092) and cultured with M-CSF (25 ng/mL) (ThermoFisher 315-02-50UG). On day 7 of culture, macrophages were polarized following existing protocols (38). For M1 macrophage polarization, LPS (InvivoGen tlrl-eblps) and IFN*γ* (Biolegend 575306) were used both at 50 ng/mL. For M2 macrophage polarization, interleukin-4 (IL4) (ThermoFisher 214-14-20UG) was used at 10 ng/mL. On day 8, the stimulation media was replaced with serum-free media (RoosterCollect™-EV, RoosterBio). Supernatant was collected and exosomes were pelleted by ultracentrifugation.

### Exosome collection

After collecting the supernatants from BMDMs and TAMs, two low-speed centrifugations (500xg) were used to remove larger unwanted particles. Then, ultracentrifugation was performed to pellet the exosomes using a Beckman Coulter Optima Max-E ultracentrifuge. Briefly, three rounds of high-speed ultracentrifuges: 20,000xg followed by two 35,000xg spins were performed. The final pellet was resuspended in 100 μL DPBS and frozen in −80 degrees Celsius.

### Exosome characterization

The Bicinchoninic Acid (BCA) Assay was used to quantify the exosomes obtained. The Nanoparticle Tracking Analyzer, Zetaview, was utilized to visualize the diameters of the exosomes.

### Sample preparation for proteomic analysis

Extracted exosomes were lyophilized by SpeedVac at 35 °C. Lyophilized exosomes were resuspended in 100 μL of 8 M urea/50 mM ammonium bicarbonate and sonicated in a bath sonicator for 30 minutes. The resulting lysates were reduced with 10 mM dithiothreitol at 37 °C for 30 minutes and subsequently alkylated with 30 mM iodoacetamide at room temperature in the dark for 15 minutes. After diluting to a final urea concentration of 1 M with 50 mM ammonium bicarbonate, sequencing-grade trypsin (Promega; Catalog #: V5111) was added at a 1:100 ratio to the total protein concentration measured. Tryptic digests were performed overnight at 37 °C and desalted via solid-phase extraction (Sep-Pak C18 1 cc Vac Cartridge, 50 mg Sorbent per Cartridge, 55-105 µm). Eluates from desalting were dried down to ∼150 μL, and 5% of each specimen was analyzed by mass spectrometry using data-dependent acquisition (DDA).

### Mass Spectrometry (MS) Analysis

MS analysis was performed in positive ion mode on a Q Exactive Plus mass spectrometer (Thermo Fisher Scientific) using a Nanospray Flex Ion Source (Cat #: ES071; Thermo Fisher Scientific). Source parameters for the DDA method are as follows: Spray voltage—2.75 kV, Capillary temperature—295°C, and S-lens RF level—50.0. Full MS parameters are as follows: Resolution—70,000, AGC target—3 × 10^6^, Maximum IT—50 ms, and Scan range—250–1,450 m/z. dd-MS^2^ parameters were as follows: Resolution—17,500, AGC target—1 × 10^5^, Maximum IT—50 ms, TopN—10, Isolation Window—1.0 m/z, and NCE—27. Data was acquired in Xcalibur (version 4.5.474.0) and searched using COMET against the reference proteome for mouse (UP000000589) from UniProt, and peptides were quantified using XPRESS. For database searching, a static modification of 57.0215 Da was added for carboxy-amido-methylation of cysteine residues, and a differential modification of 15.9949 Da was added for oxidized methionine residues. The search results were manually inspected and filtered so all identifications satisfy the following criteria: mass difference within ± 15 ppm of the calculated peptide mass, XPRESS probability greater than 0.7, and unique to a single protein in the database.

### Data analysis

#### Sample selection and filtering

In total, 3 MS studies were performed on 30 macrophage EV samples (12 were from BMDMs (4 M0, 5 M1, 3 M2), 18 were from TAMs (5 hMEK, 7 control/parent)). The first MS contained only BMDMs EVs, while the other two contained both TAMs EVs and control BMDMs EVs. For further analysis to determine characteristic proteins, all 12 BMDMs-EVs were examined and 13 out of 18 TAMs-EVs that have more than 8 average reliable proteins were investigated. Therefore, 25 EV samples proceeded to downstream analysis.

#### Unsupervised hierarchical clustering

In total, 131 and 138 proteins were detected in the second and third experiments, respectively. Out of these proteins, 55 proteins were detected in both MS experiments. 2 EV samples from M1 BMDMs contained no expression of any of the 55 common proteins, and are therefore excluded from downstream analysis, resulting in 23 EV samples. 74 proteins were detected in the first MS, and 12 out of these 74 were shared with the 55 common proteins and therefore are conserved across all three MS. To ensure these protein expression trends can provide accurate reflections of EV patterns, we removed 12 proteins, including keratin, that are known to be contaminants in MS (39). To examine how the 23 macrophage EV samples express the remaining 43 proteins, we utilized Morpheus (https://software.broadinstitute.org/morpheus) to generate a Kendall similarity matrix based on the number of unique peptides for each protein contained within each sample and performed a Pearson’s unsupervised hierarchical clustering.

#### Signature EVs proteins for M1/M2 BMDMs

We identified 17 and 42 signature EV proteins for M1 and M2 BMDMs, respectively. The steps for pinpointing these proteins are included in Supplemental Figure 3. The identities of these proteins are reported in Supplemental Table 1, in both Uniprot protein IDs and actual protein names (40). Briefly, for each of the 3 MS experiments, we overlayed the proteins found in M1 BMDMs EVs with those found in M2 BMDMs EVs and presented them using Venn diagrams tools in Hiplot, (ORG) (https://hiplot.org), a comprehensive and easy-to-use web service for boosting the publication-ready biomedical data visualization. Removing the proteins that are shared, we focused on the proteins that are unique to M1 and M2 BMDMs EVs. Across the three MS, we assembled the proteins unique to M1 BMDMs EVs together and acquired a list of 17 proteins; we did the same procedure for M2 BMDMs EVs and acquired a list of 42 proteins. Apart from our data, there are two other published works with M1/M2 BMDMs EVs signature proteins (35, 41).

#### Signature EV proteins for regressor and progressor TAM EVs

For EV proteins from regressor and progressor TAMs, we first prepared a list of proteins that are expressed at least once in the regressor TAM-EVs from two MS experiments. We then overlapped the two lists to acquire a list of EVs proteins that are expressed at least once in regressor-TAM-EVs but are conserved between the two MS experiments. The same steps were performed for progressor-TAM-EVs. The two filtered protein lists were treated as signature proteins for regressor- and progressor-TAM-EVs. We then overlapped the two lists to remove the proteins that are common to both progressor- and regressor-TAM-EVs, thus looking at the proteins that are truly unique. We then overlapped the proteins in this new list with M1 and M2 EV signature proteins.

#### PPI networks and Gene Ontology (GO) enrichment for biological processes

Protein-protein interaction networks were performed utilizing STRING, version 12.0 (42). The two protein lists generated in the determination of signature EV proteins for progressor- and regressor-TAM-EVs were utilized. Gene Ontology (GO) enrichment analysis was performed to determine the enriched biological processes in the EVs, with pathways with FDR ≤ 0.05 considered to be significant.

## Results

### Isolation of TAMs from mouse tumors and purification of TAM-EVs

To elucidate the protein cargo in EVs from TAMs, we isolated TAMs directly from mouse 9609 progressor and regressor tumors, cultured them in serum-free media for either short-term or long-term, and purified their EVs from supernatants. We used magnetic bead isolation to positively select F4/80+ cells from a tumor single-cell suspension and assessed their purity by measuring their CD11b surface expression by flow cytometry (Figure 1A) across 12 samples (Figure 1B). Study of the expression of MHC class II as a marker of M1-like polarization (37, 43) indicated that, consistent with previous studies, TAMs purified from regressors have higher average MHC class II expression than the TAMs acquired from progressors (Figure 1E) (36, 37).

**Figure 1.**
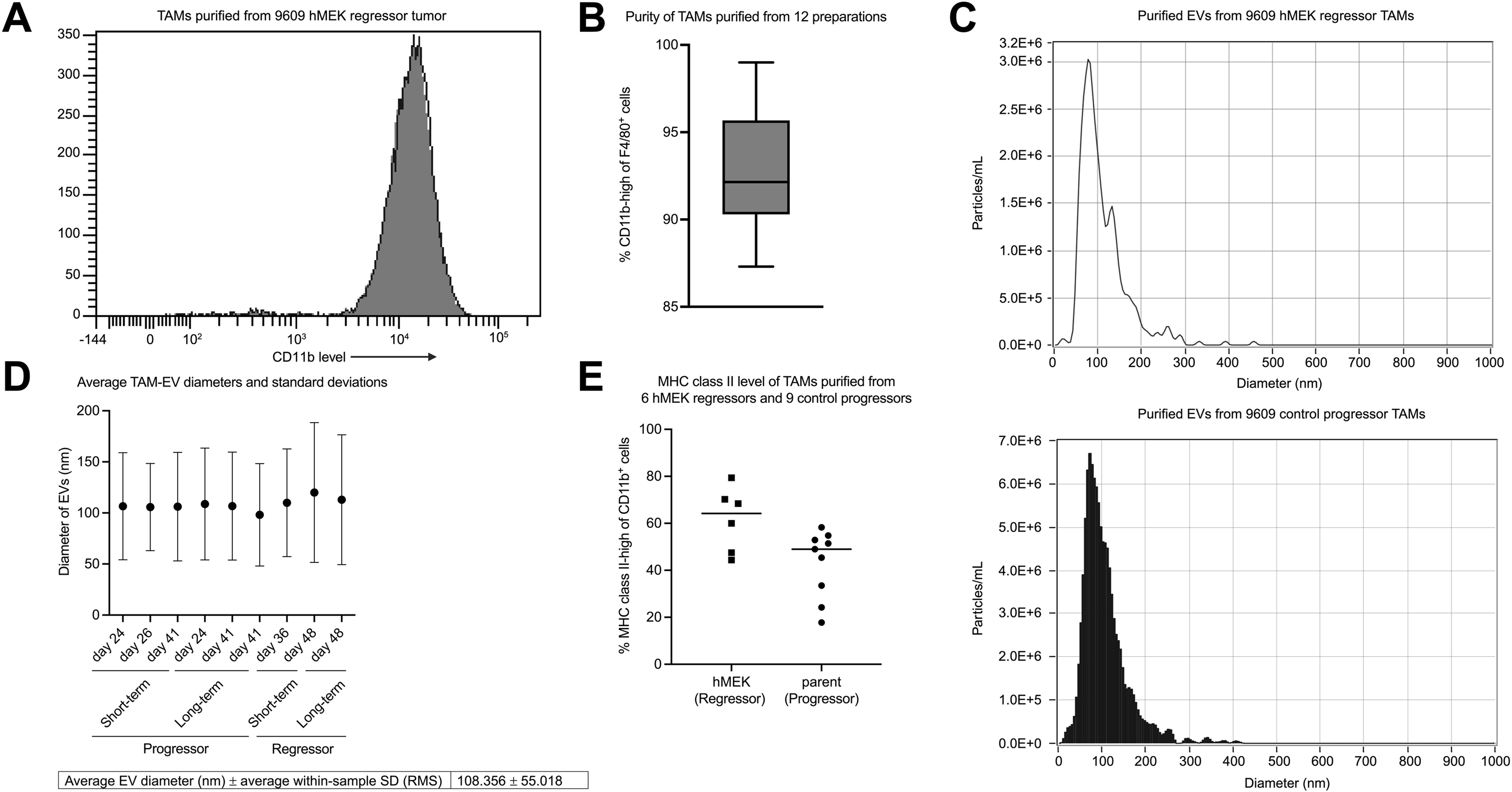
Characterization of TAM and EV preparations. A: FACs staining for CD11b on F4/80+ cells to determine TAM purity for one TAM sample. B: The overall TAM purity from 12 preparations, average purity = 93.0%. C: NTA Zetaview plot for EVs from a regressor (top) and a progressor (bottom) TAM-EV sample. D: EV average diameter and within-sample standard deviation from 9 preparations, calculated using root mean square equation. The table reports the unweighted average of the mean EV diameters and the average within-sample standard deviation. E: MHC class II level of TAMs purified from progressor or regressor tumors.

Next, we purified EVs from TAM supernatants and determined their purity using Zetaview Nanoparticular Tracking Analyzer (NTA). Figure 1C shows an example plot of EVs purified from regressor and progressor TAMs, demonstrating that most of the particles fell within the exosome size range of 30-150 nm (44), with a mean diameter of 108.356 nm and standard deviation of 55.018 nm across nine random preparations (Figure 1D).

### Long-term culture alters the protein profile of TAM-EVs

Having successfully purified TAMs and their EVs, we next sought to examine the protein cargo in TAM-EVs compared to EVs from BMDMs. Three MS studies were performed on 30 macrophage-derived EV samples (12 from BMDMs and 18 from TAMs). The full workflow for the filtering and selection of samples to analyze is detailed in the methods section. Briefly, the original 30 samples were filtered into the final 23 based on having EVs that contained more than 8 proteins and also had protein identities found in the the 43 conserved proteins. Among EV proteins, 55 were found to be conserved across multiple MS studies and 43 of these were determined to be non-contaminants. Of note, the number of proteins detected in EVs was not dependent on the concentration of the proteins in the EVs (Supplemental Figure 1A). Venn diagram analyses of the protein content revealed substantial overlap among replicate samples, whereas no obvious shared proteins were observed across different samples (Supplemental Figure 1B and 1C).

To get insight into the relatedness of EVs purified from different macrophage sources, unsupervised hierarchical clustering was applied on the 23 EV samples based on the 43 conserved proteins, utilizing the metric of one minus Pearson correlation (Figure 2A). Overall, the analysis demonstrated three main findings: 1) EVs from biological replicates clustered together, confirming the reliability of our approach to harvest EVs from TAMs; 2) EVs from TAMs clustered together based on the duration of in vitro culture, rather than on how long the tumor had been growing in the mouse or its phenotype (progressor or regressor); 3) EVs from long-term cultured TAMs tended to cluster with EVs from BMDMs. For example, EVs from regressor (d48) TAMs cultured long-term clustered more closely to the EVs from progressor (d24) TAMs cultured long-term, rather than the other two samples also from regressor tumors that underwent short-term culture. These results suggest that prolonged in vitro culture can reshape EV protein cargo and also demonstrate that our EV isolation and MS protocol can give rise to interpretable data and provide logical and reproducible clustering findings.

**Figure 2.**
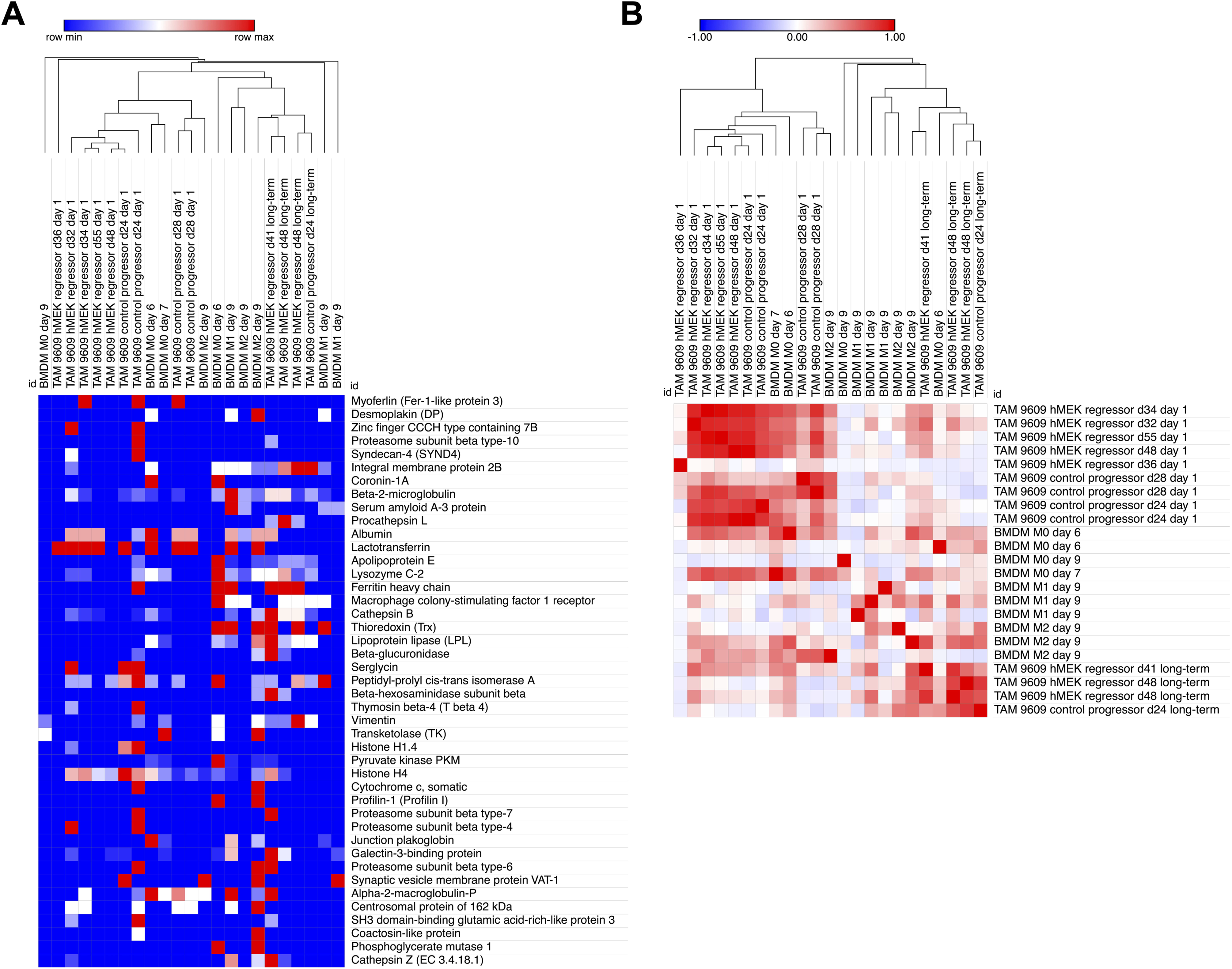
Comparison of EV proteins across all samples shows in vitro culture determines relatedness of the EV protein content. A: Unsupervised hierarchical clustering of 23 samples based on 43 conserved proteins. Samples labeled by source of macrophage (BMDM vs TAM), day of tumor harvest (for TAMs only), phenotype of tumor (for TAMs only), and duration of culture in vitro. The one minus Pearson correlation metric was performed. B: Unsupervised hierarchical clustering performed on the similarity matrix comparing the 23 EV samples to each other based on their protein expressions, using Kendall metric.

To corroborate this observation, we next generated a similarity matrix, where each EV sample is compared to the other 22 samples based on their expression of the 43 conserved EV proteins (Figure 2B). The unsupervised hierarchical clustering performed on this similarity matrix recapitulated the trends observed in Figure 2A. Notably, the clustering clearly separated the TAM-EV samples into two major groups: EVs from TAMs after a 1-day culture and EVs from TAMs cultured for more than 1 day, highlighting the impact of culture duration on EV protein cargo. The EVs from TAMs derived from a day 48 (d48) 9609 hMEK regressor tumor is of particular interest. In this experiment, EVs were harvested from the exact same TAMs after short-term (day 1) or long-term (day 5 and day 7) in vitro culture. In the clustering, the EVs from the exact same TAMs were not clustered together, but rather clustered with EVs from other TAMs based on duration of culture. This observation is also supported by the analysis of EV samples from a day 24 (d24) 9609 control progressor TAMs, which again clustered according to time in culture rather than source of macrophage.

Interestingly, EVs from BMDMs also demonstrated clustering tendencies with EVs from long-term and short-term plated TAMs. EVs from 9-day cultured BMDMs (M0, M1, and M2) clustered with EVs from long-term cultured TAMs, whereas EVs from 6- or 7-day cultured BMDMs (the minimum culture duration to differentiate macrophages from hematopoietic stem cells) clustered with EVs from short-term cultured TAMs, suggesting that extended in vitro culture might change the protein cargo of EVs. This suggests that in vitro culture created similarities in EV cargo regardless of polarization state or source of the macrophage.

### Prolonged in vitro culture shifts TAM-EV protein content towards BMDM-EVs

Through unsupervised hierarchical clustering, we observed that EVs from long-term in vitro culture of TAMs clustered distinctly from EVs from short-term culture of TAMs. Thus, we investigated the effects that long-term plating might have on the TAMs themselves. We found that over the course of 7 days, MHC class II levels gradually diminished on TAMs from a 9609-hMEK (regressor) tumor (Figure 3A-B). This gradual loss of MHC class II also occurred with 3 other regressor-TAMs and 3 progressor-TAMs, even though progressor-TAMs initially exhibited lower overall MHC class II levels on day 0, compared to their regressor counterparts (Figure 3C).

**Figure 3.**
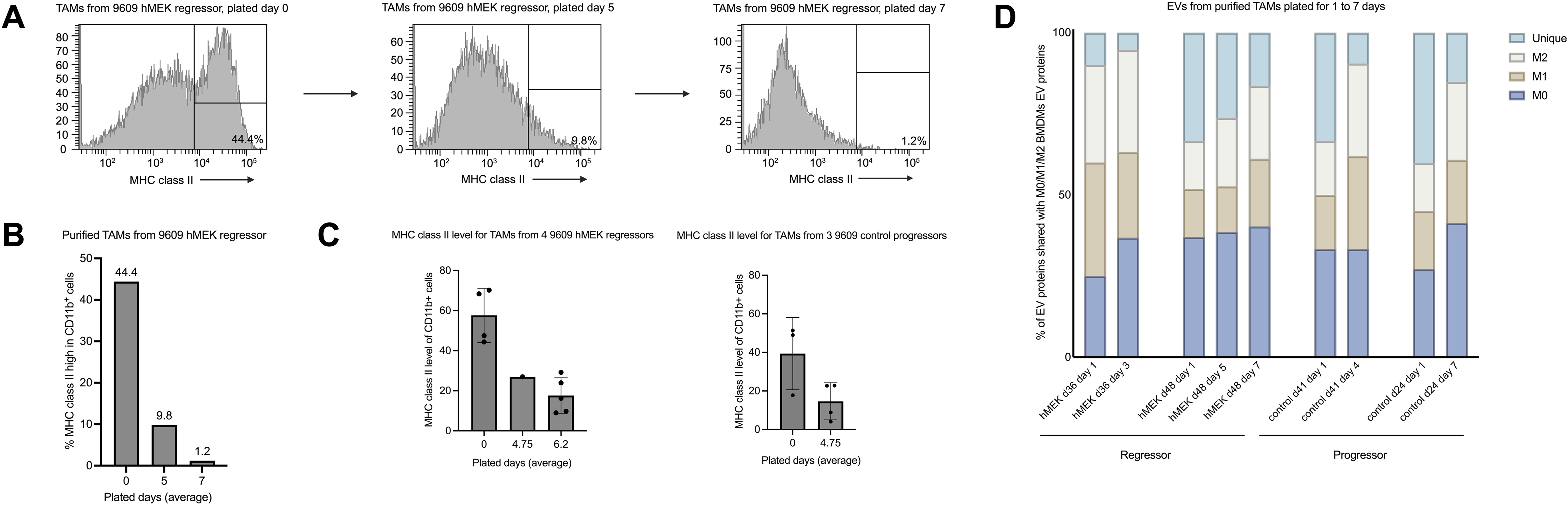
In vitro culture of TAMs changes their phenotype and reduces unique proteins in their EVs. A: FACs staining of MHC class II levels on TAMs isolated from 9609-hMEK1 tumors over the course of 7 days of in vitro culture. B: Statistical presentations of trends in Figure 3A. C: The overall change in MHC class II from day 0 to a long-term plated day, comparing regressor-TAMs and progressor-TAMs. D: EVs collected from TAMs on day 0 and long-term plated day(s), comparing the EV proteins in the specific TAMs with the proteins found in M0/M1/M2 BMDMs-EVs.

Next, we asked how long-term plating of TAMs affected the protein cargo in the EVs. To capture temporal changes in TAM-EV protein cargo, we collected EVs sequentially from the same TAM cultures at defined time points and analyzed the percentage of protein shared with M0/M1/M2 BMDM-EVs. We noticed that long-term plating caused a gradual decrease in proteins unique to TAM-EVs and a gradual increase in proteins shared with M0 and M2 BMDM-EVs (Figure 3D). These data indicate that long-term culture of TAMs leads to a loss or alteration of their differentiated phenotype, producing EVs that carry more “generic” cargo rather than unique proteins.

### Identification of protein signatures associated with short-term cultured progressor and regressor TAM-EVs

Having shown that long-term culture of TAMs might change their differentiation state and cause reductions in unique proteins in the EVs, we next studied the cargo in EVs from short-term plated TAMs, comparing it with cargo in EVs from BMDMs. We hypothesized that EVs from M1 BMDMs would have proteins similar to EVs from regressor-TAMs; however, we found that total proteins in EVs from M1 or M2 BMDMs did not preferentially match with those from TAM-EVs, regardless of regressor or progressor origin or how long the tumor had been growing in the mouse (Supplemental Figure 2). Since this global comparison did not specifically identify signature proteins, we next sought to restrict the comparison to unique proteins in the EVs from short-term TAMs versus M1 and M2 BMDMs. We first identified EV protein signatures for M1 and M2 BMDMs (Supplemental Figure 3) and compared them with two previous studies (35, 41). Surprisingly, little similarity was found between protein content of BMDM-EVs across all three studies (including ours), suggesting lab-to-lab variability. One similarity, however, was that both Cianciaruso et al.’s and our study reported fewer signature EV proteins from M1 than M2 BMDMs, suggesting that M1 BMDM-EVs contained a lower diversity of proteins.

Next, we set out to identify the EV protein signature of progressor- and regressor-TAMs. Figure 4A shows the workflow in which 11 EV samples from short-term cultured TAMs (5 from regressor and 6 from progressor TAMs) were evaluated separately. Based on the protein expression in at least one sample, a list of 41 and 56 proteins were identified in regressor- and progressor-TAM-EVs respectively. After removal of contaminant proteins, we then overlapped the two lists from the two MS experiments in both progressor- and regressor-TAM-EVs, identifying 4 and 6 signature proteins in regressor TAM-EVs and progressor TAM-EVs, respectively.

**Figure 4.**
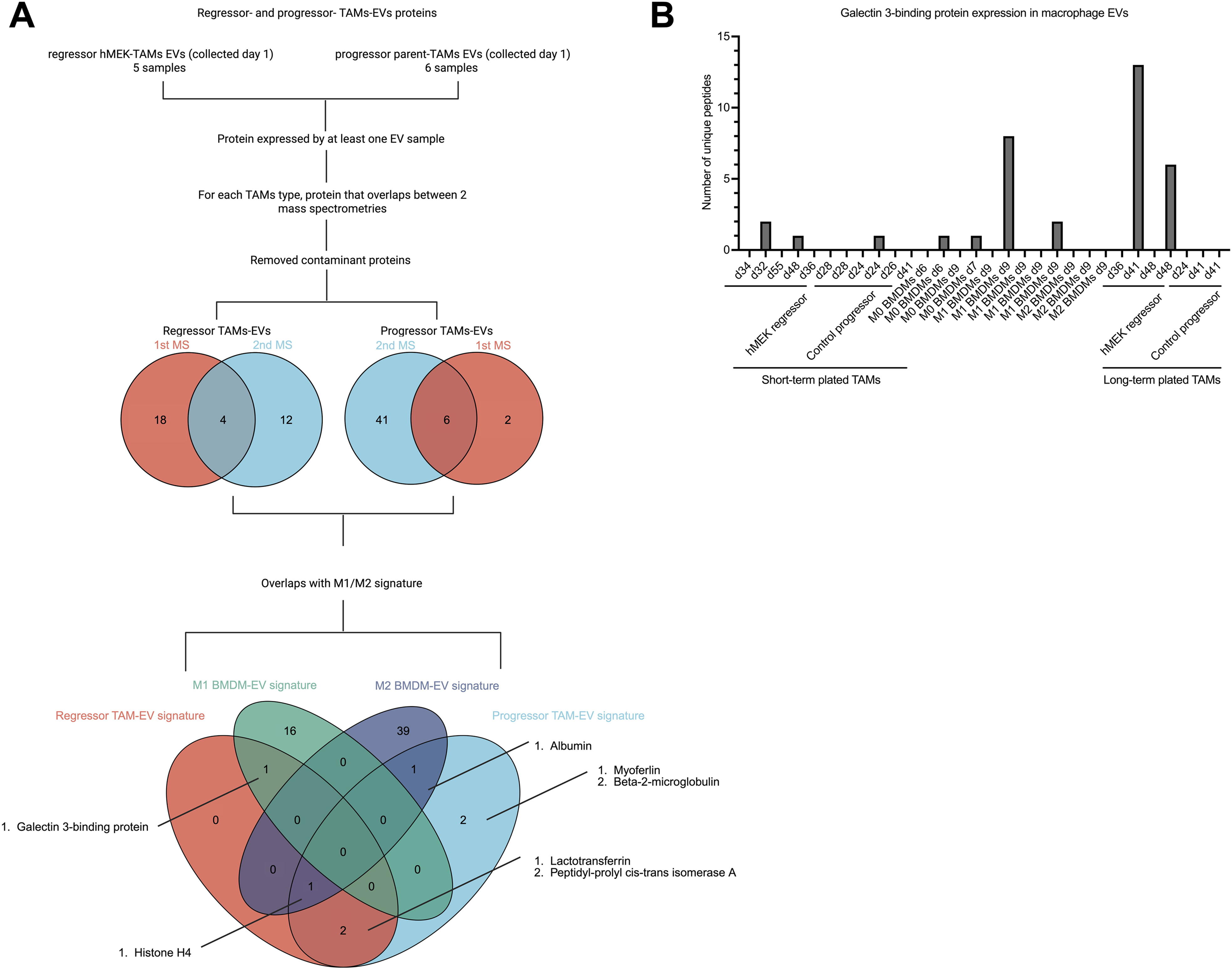
Identification and comparison of signature proteins in TAM and BMDM EVs. A: Workflow for determination of EV proteins that are shared or unique between progressor-and regressor-TAMs, and also shared with M1 and M2 BMDM EV signature proteins (Supplemental Figure 3 and Supplemental Table 1). B: Macrophage EV samples that contain galectin 3-binding protein (LGALs3bp), presenting the number of unique LGALs3bp peptides detected by MS in the 30 total EV samples.

We compared these regressor-and progressor-TAM-EV protein signatures with the signature proteins we had identified in EVs from M1 and M2 BMDMs EV (Supplemental Figure 3 and Supplemental Table 1) and present the results in the 4-element Venn diagram (Figure 4A). The results indicate that of the 4 signature proteins from regressor TAM-EVs, 1 overlapped with proteins from M1-EVs, 1 overlapped with proteins from both M2-EVs and progressor TAM-EVs, and 2 overlapped only with progressor TAM-EVs. Of the 6 signature proteins from progressor TAM-EVs, 1 overlapped with proteins from M2-EVs, 1 overlapped with proteins from both M2-EVs and regressor TAM-EVs, 2 overlapped with regressor TAM-EVs, and 2 were unique.

We next focused on galectin 3-binding protein because it was common between regressor TAM-EVs and M1 BMDM-EVs. Examining the original 30 samples, we detected this protein in all three MS experiments. 9 out of the 30 EV samples expressed galectin 3-binding protein, 4 from regressor TAM-EVs, 2 from M0 BMDM-EVs, 2 from M1 BMDM-EVs and 1 from progressor TAM-EVs. Interestingly, this protein was completely absent in all the M2 BMDM-EVs and 4 of the 5 progressor TAM-EV samples.

### EVs from short-term cultured progressor TAMs are enriched in immunostimulatory pathways

To determine pathways that may mediate functional outcomes of TAM-derived EVs, we performed a Protein-Protein Interaction (PPI) analysis of the proteins found in EVs from short-term plated TAMs. We first compiled the 56 and 41 proteins previously identified as the signatures for progressor and regressor TAM-EV samples found in the previous result section (Figure 4A). The PPI analysis of these 56 and 41 proteins showed a prominent cluster that included different versions of keratin (Figures 5A-B). This keratin cluster was connected to the rest of the groups by fewer lines. This observation, along with previous publications identifying keratin as a contaminant protein in MS (39, 45), encouraged us to remove these proteins and repeat the PPI study in a filtered subset of proteins (Figure 5C and D).

**Figure 5.**
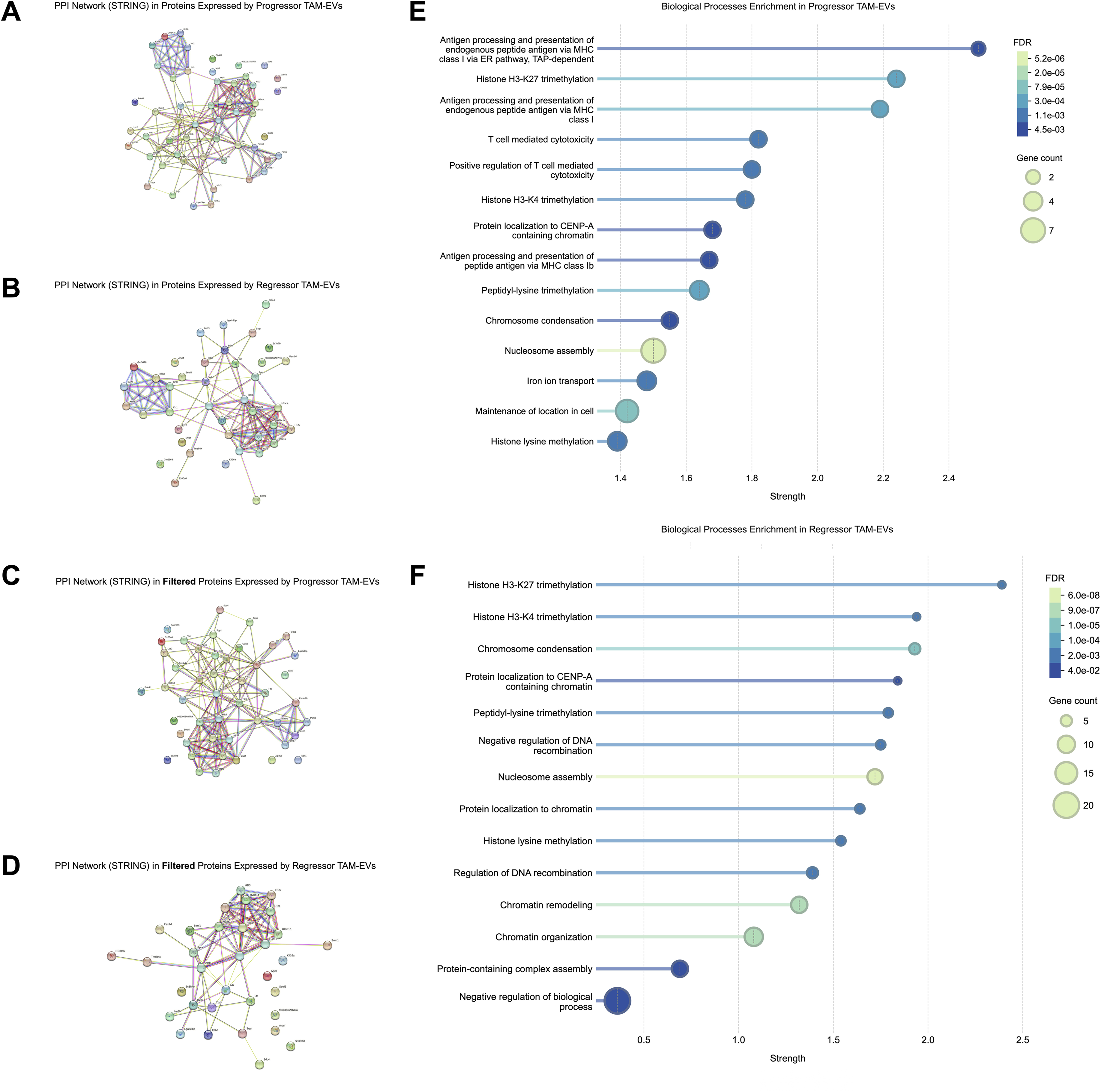
Protein pathways in TAM-EVs include immune activation networks in EVs from progressor-TAMs. PPI networks generated by STRING, with the association diagrams focusing on proteins that are expressed by at least one sample in A: progressor-TAM-EVs and B: regressor-TAM-EVs. The proteins were then filtered to remove proteins known to be the contaminants in MS, thereby generating networks with filtered proteins in C: progressor-TAM-EVs and D: regressor-TAM-EVs. Top biological pathways are shown in E: progressor-TAM-EVs and F: regressor-TAM-EVs.

From the PPI network of filtered proteins, we list the top 14 pathways with the greatest strength (confidence score) for both progressor-TAM-EVs (Figure 5E) and regressor-TAM-EVs (Figure 5F). Interestingly, pathways enriched in progressor-TAM-EVs uniquely included many pathways engaged in immune stimulation. Two of the top three pathways in EVs from progressor-TAMs are involved in antigen presentation via MHC class I, with the next two pathways participating in T cell cytotoxicity. These pathways were not present in regressor-TAM-EVs, in which the primary pathways were those that participate in epigenetic processes (e.g. methylation, chromatin dynamics and nucleosome organization).

## Discussion

Previously, we found that EVs from regressor tumor cells could change macrophage gene expression due to the expression of the protein MEK1, which was enriched in EVs from regressor compared to progressor tumor cell lines (36). This finding encouraged us to hypothesize that protein content in EVs could indeed alter target cell activity. In this study, we characterized the protein content of 30 independent preparations of EVs released by macrophages, including 18 TAMs and 12 BMDMs. We analyzed EVs from TAMs purified from both progressor and regressor tumors and compared them with EVs from BMDMs. In contrast to previous studies that assessed EVs from progressor TAMs via sorting of EVs and a subtractive approach (35), we purified TAMs (>90% purity) directly from both regressor and progressor tumors and harvested EVs via ultracentrifugation for MS studies. Our results highlight a feasible protocol to study EVs from macrophages and identify specific proteins that might be unique in regressor versus progressor TAMs. Surprisingly, we found that EVs from progressor TAMs might have greater immune stimulatory proteins than EVs from regressor TAMs, but the functional consequence of this protein difference is not known. Given the small quantities of EVs we could harvest, functional studies will be performed in future investigations.

Through unsupervised hierarchical clustering of data obtained from MS studies of EV preparations, we observed that replicate EVs clustered together and moreover, EVs from TAMs clustered based on duration of culture in vitro, rather than origin in vivo. This finding validated that our approach could provide consistent and interpretable results. Even though the “ground truth” of which EVs should cluster together is not known, our methods grouped EV replicates together. Thus, we conclude based on this analysis that 1) the cargo of EVs from macrophages reflects the duration of in vitro culture, and 2) the polarization state of the macrophages, i.e., M0, M1, M2, regressor-TAM, progressor-TAM, had little effect on overall protein content and identity. Notably, we found that long-term in vitro plating reduced MHC class II levels in TAMs, while the corresponding EVs from the TAMs showed increased proteins shared with M0 and M2 BMDM-EVs and decreased proteins unique to TAM-EVs. These results suggest that the differentiation or polarization state of TAMs could be lost or altered over time and highlight the importance of studying TAMs immediately after harvest to preserve their functional status in vivo. In fact, two studies have documented the effects of extended vitro culture on macrophages. Subramanian et al. demonstrated that alveolar macrophages, after 2 months of ex vivo expansion, had significant alterations in their transcriptome but restored their epigenetic identities after being re-introduced into their in vivo niche (46). Chamberlain et al. reported that 21 days of in vitro culture of primary BMDMs and macrophage cell lines conferred a more M2-like phenotype (47).

Focusing on short-term plated TAMs, we identified signature proteins from EVs isolated from progressor- and regressor-TAMs and compared them with EV signature proteins from M1 and M2 BMDMs. We found galectin 3-binding protein (LGALs3bp) to be shared between regressor TAM-EVs and M1 BMDM-EVs. LGALs3bp is a glycoprotein that has both intracellular and extracellular actions, including increasing angiogenesis, enhancing tumor invasion, and repressing fibrocyte differentiation (48–50). When secreted, LGALs3bp could be released in a free soluble form, or bound to EVs (48, 51). Past studies have shown LGALs3bp to be enriched in the tumor microenvironment, and that its major source was tumor cells (52, 53). Given the pro-tumor effects of LGALs3bp, our results showing that LGALs3bp is found in EVs from pro-inflammatory and anti-tumor macrophages suggest possible anti-tumor abilities. In fact, several studies have also demonstrated the potential anti-tumor activities of LGALs3bp in colorectal cancer, where it suppresses WNT signaling and could promote a good prognosis (54, 55). The main binding partner of LGALs3bp is galectin-3 (gal-3), but studies have uncovered other binding partners such as galectin-1 (gal-1) (56, 57). Interestingly, both gal-1 and gal-3 were identified in an EV sample from M2 BMDMs. The presence of LGALs3bp in EVs from pro-inflammatory and anti-tumor macrophages and its binding partners in EVs from anti-inflammatory macrophages may imply some level of crosstalk between different types of macrophages, although more studies should be done to confirm this pattern.

Previous studies found that EVs from macrophages had distinct protein content from the cytoplasmic proteins and even suggested that EVs promote distinct biologic activities compared to their parent cells (35, 41). We therefore identified protein pathways that could be enriched in EVs from macrophages and found that surprisingly EVs from progressor-TAMs possess greater immunostimulatory protein pathways than EVs from regressor-TAMs. Both the progressive growth pattern of the progressor tumors and the lower MHC class II levels of progressor-TAMs led to the expectation that their EVs contain immunosuppressive and not stimulatory proteins. As antigen-presenting cells, macrophages utilize MHC class II to present extracellular peptides to activate CD4+ T cells. (58, 59). Macrophages can also activate CD8^+^ T cells through cross-presentation of extracellular peptides or endogenous presentation of intracellular peptides on MHC class I (60, 61). We found enrichment of the MHC class I antigen presentation pathway and two T cell cytotoxicity pathways in EVs from progressor-TAMs versus regressor-TAMs. These results contrast with our previous findings that regressor tumors had more cytotoxic lymphocytes than progressor tumors (36, 37). Future studies will investigate the target tissues that uptake EVs from regressor-versus progressor-TAMs. We speculate that EVs from progressor-TAMs may seed distant organs and stimulate inflammation therein to facilitate tumor metastasis, whereas EVs from regressor-TAMs might reduce inflammation systemically. Although tumor-derived EVs have been tracked and shown to seed metastatic sites (62, 63), whether EVs from TAMs reach similar distal organs has not been shown to our knowledge.

When analyzing the MS data, we noticed that the protein cargo, even those among biological replicates, had variability (Supplemental Figure 1). This is likely due to the fact that the EVs themselves possess heterogeneity, even when released by the same cell (64, 65). We imagine that the diversity in EV cargo may be even greater if the cells, e.g. TAMs, are purified from tumors on different days from different mice. Although our ultracentrifugation method of purification could also induce heterogeneity, other EV purification methods likely introduce even more variability (66). Nevertheless, we found that technical triplicates showed high overlap of protein identities (Supplemental Figure 1C), suggesting relatively low variabilities produced by ultracentrifugation and MS procedures. This observation encouraged us to regard EVs’ own heterogeneity as the main contributor to variability. Indeed, when we compared the protein identities in our EVs with two previous studies (Supplemental Figure 3), there was little overlap between the protein cargo across the three studies, suggesting high lab-to-lab variability. Overall, the trends we identified in this report provide proof-of-concept that EVs from macrophages indeed have distinct cargo that could reflect in vivo biology, thus propelling enthusiasm for future functional studies. The inherent biologic variability of EVs and TAMs are limitations of these studies, and standardization of TAM isolation, TAM culture, and EV harvest will benefit the field and allow for future functional studies to yield significant and reproducible findings.

## Acknowledgment

The work was supported by Office of Research Infrastructure Programs of the National Institutes of Health under award number 1S10OD036412-01. We thank openbiox community and Hiplot team (https://hiplot.org) for providing technical assistance and valuable tools for data analysis and visualization.

## Funding

This project was supported by a grant from The Hartwell Foundation awarded to J.D.B. This project was also supported by a NICHD grant (1UG1HD119608) awarded to R.T.S.

## Conflict of Interest Statement

The authors declare no conflicts of interest.

## Author contribution

**Yining Zhu:** Conceptualization (supporting); data curation (lead); formal analysis (lead); investigation (lead); methodology (lead); visualization (lead); writing—original draft (lead); writing—review and editing (supporting). **Jiaxi Cai:** Data curation (supporting); investigation (supporting); visualization (supporting); writing – review and editing (supporting). **Raymond T. Suhandynata:** Funding acquisition (supporting); project administration (supporting); visualization (supporting); writing—review and editing (supporting). **Jack D. Bui**: Conceptualization (lead); data curation (supporting); formal analysis (supporting); funding acquisition (lead); investigation (supporting); methodology (supporting); project administration (lead); resources (lead); supervision (lead); validation (supporting); visualization (supporting); writing—original draft (supporting); writing—review and editing (lead).

## Notes

### Competing Interest Statement

The authors have declared no competing interest.

