## Supplementary figures and images for "Identification Of Protein Cargo in Extracellular Vesicles from Macrophages in Progressing and Regressing Tumors"

### Supplemental Figure 1

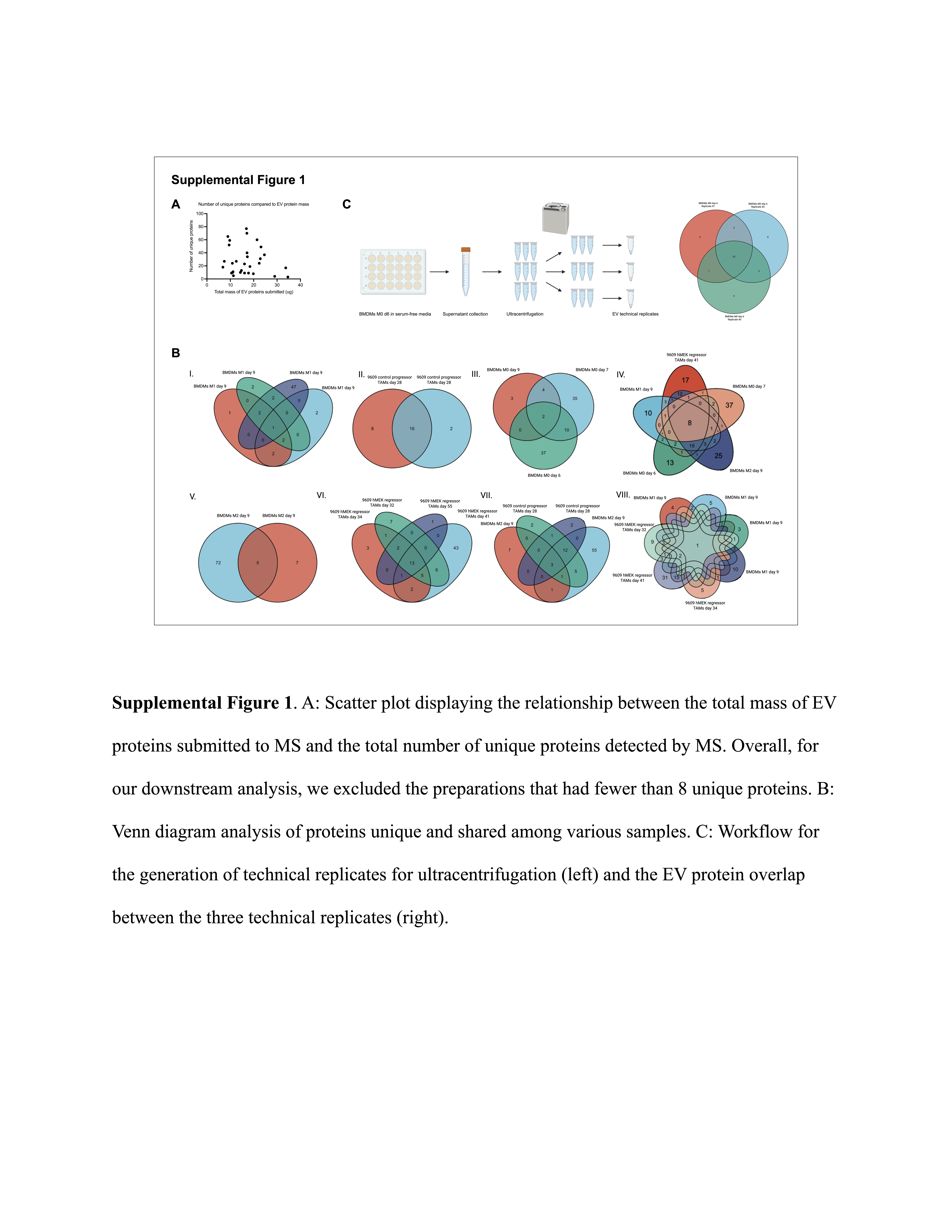

### Supplemental Figure 2

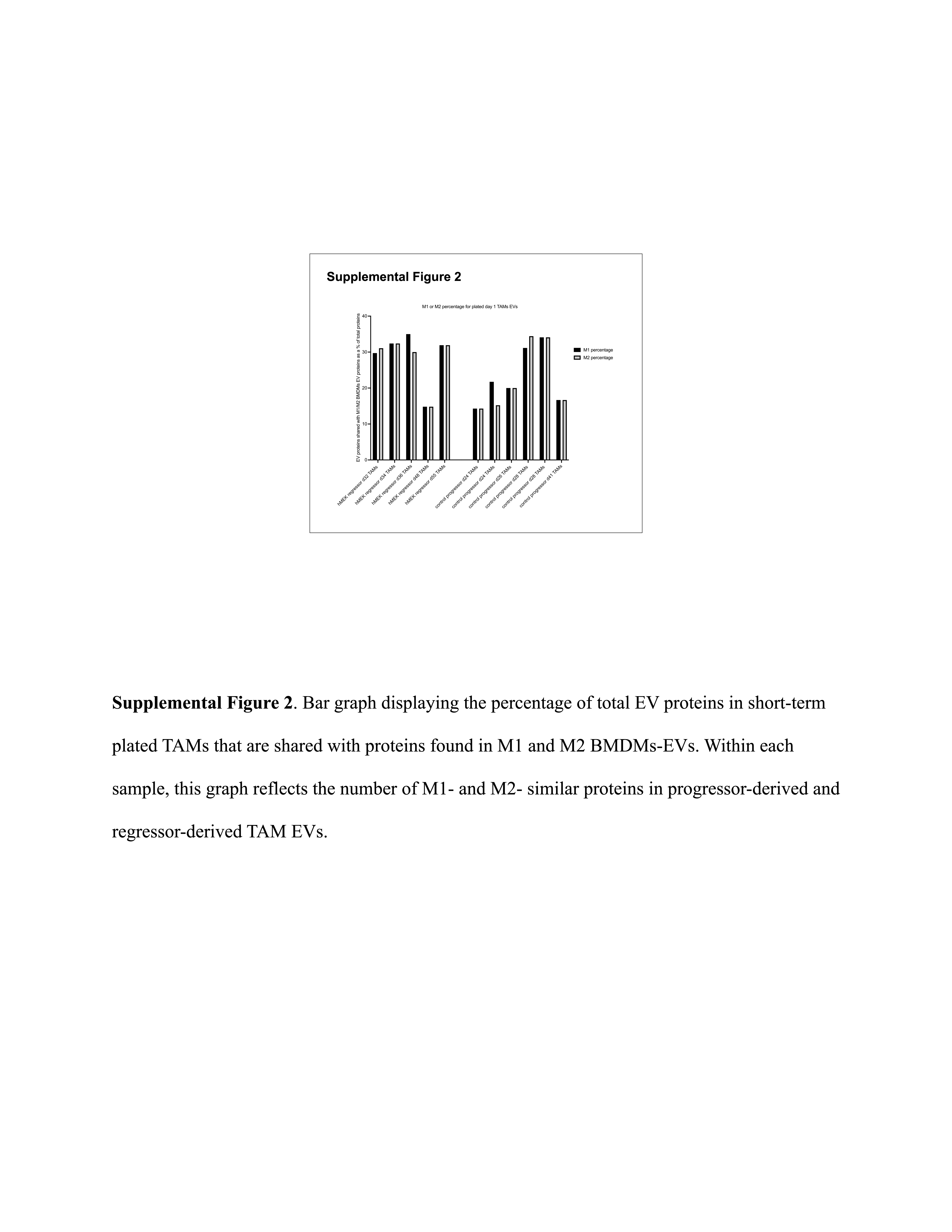

### Supplemental Figure 3

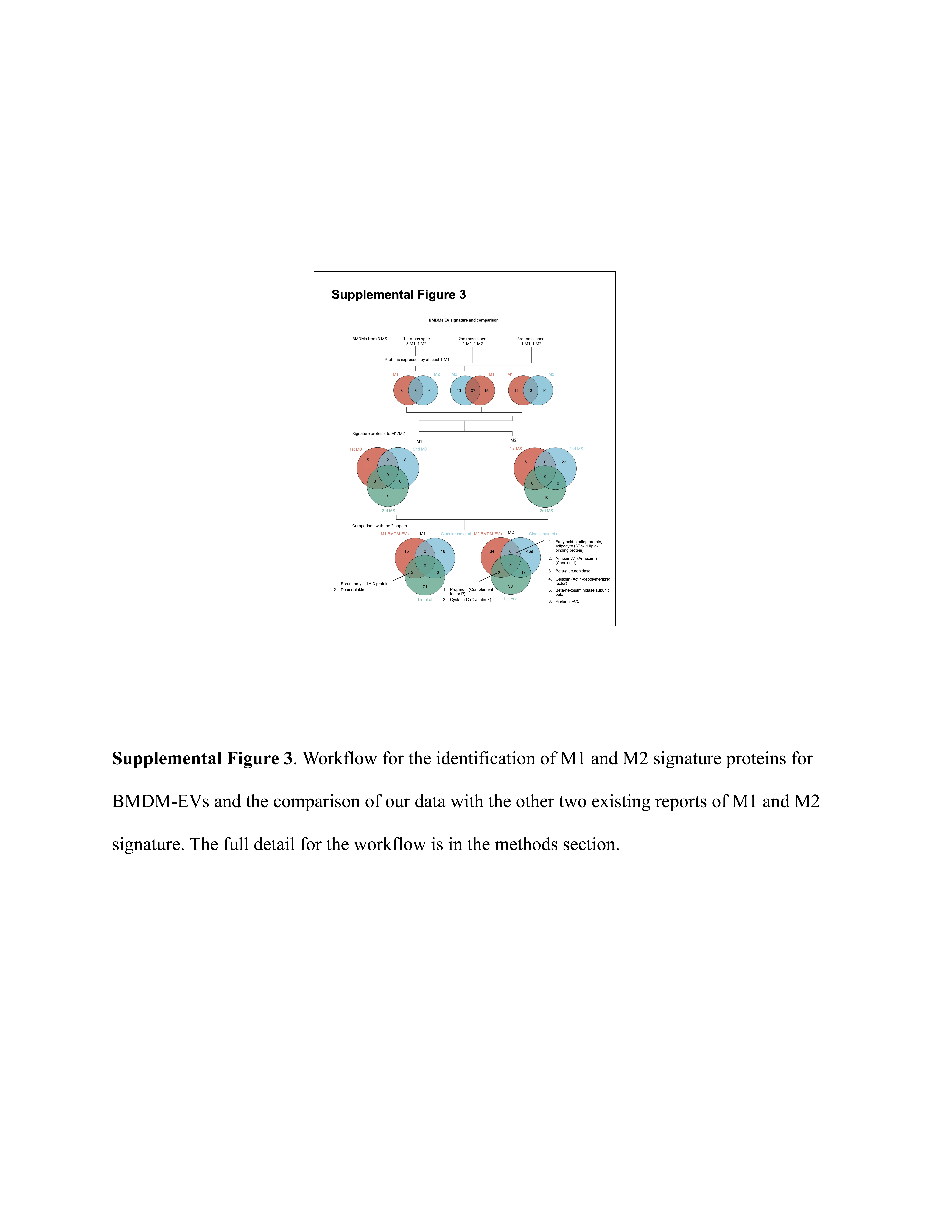
