## Supplemental Table 1 for "Identification Of Protein Cargo in Extracellular Vesicles from Macrophages in Progressing and Regressing Tumors"

**M1 BMDM-EV signature proteins**

| From | Entry | Reviewed | Entry Name | Protein names | Gene Names | Organism | Length |
|---|---|---|---|---|---|---|---|
| P04918 | P04918 | reviewed | SAA3_MOUSE | Serum amyloid A-3 protein | Saa3 | Mus musculus (Mouse) | 122 |
| P17182 | P17182 | reviewed | ENOA_MOUSE | Alpha-enolase (EC 4.2.1.11) (2-phospho-D-glycerate hydro-lyase) (Enolase 1) (Non-neural enolase) (NNE) | Eno1 Eno-1 | Mus musculus (Mouse) | 434 |
| Q07797 | Q07797 | reviewed | LG3BP_MOUSE | Galectin-3-binding protein (Cyp-C-associated protein) (CyCAP) (Lectin galactoside-binding soluble 3-binding protein) (Protein MAMA) | Lgals3bp Cycap Mama | Mus musculus (Mouse) | 577 |
| Q61207 | Q61207 | reviewed | SAP_MOUSE | Prosaposin (Sulfated glycoprotein 1) (SGP-1) [Cleaved into: Saposin-A; Saposin-B-Val; Saposin-B; Saposin-C; Saposin-D] | Psap Sgp1 | Mus musculus (Mouse) | 557 |
| A0A0U1RNR0 | A0A0U1RNR0 | unreviewed | A0A0U1RNR0_MOUSE | Hormone-sensitive lipase (EC 3.1.1.23) (EC 3.1.1.79) (Monoacylglycerol lipase LIPE) (Retinyl ester hydrolase) | Lipe | Mus musculus (Mouse) | 180 |
| P01027 | P01027 | reviewed | CO3_MOUSE | Complement C3 (HSE-MSF) [Cleaved into: Complement C3 beta chain; C3-beta-c (C3bc); Complement C3 alpha chain; C3a anaphylatoxin; Acylation stimulating protein (ASP) (C3adesArg); Complement C3b (Complement C3b-alpha' chain); Complemen | C3 | Mus musculus (Mouse) | 1663 |
| P09528 | P09528 | reviewed | FRIH_MOUSE | Ferritin heavy chain (Ferritin H subunit) (EC 1.16.3.1) [Cleaved into: Ferritin heavy chain, N-terminally processed] | Fth1 Fth | Mus musculus (Mouse) | 182 |
| P09581 | P09581 | reviewed | CSF1R_MOUSE | Macrophage colony-stimulating factor 1 receptor (CSF-1 receptor) (CSF-1-R) (CSF-1R) (M-CSF-R) (EC 2.7.10.1) (Proto-oncogene c-Fms) (CD antigen CD115) | Csf1r Csfmr Fms | Mus musculus (Mouse) | 977 |
| P14106 | P14106 | reviewed | C1QB_MOUSE | Complement C1q subcomponent subunit B | C1qb | Mus musculus (Mouse) | 253 |
| P98086 | P98086 | reviewed | C1QA_MOUSE | Complement C1q subcomponent subunit A | C1qa | Mus musculus (Mouse) | 245 |
| P98203 | P98203 | reviewed | ARVC_MOUSE | Splicing regulator ARVCF (Armadillo repeat protein deleted in velo-cardio-facial syndrome homolog) | Arvcf | Mus musculus (Mouse) | 962 |
| Q60963 | Q60963 | reviewed | PAFA_MOUSE | Platelet-activating factor acetylhydrolase (PAF acetylhydrolase) (EC 3.1.1.47) (1-alkyl-2-acetylglycerophosphocholine esterase) (2-acetyl-1-alkylglycerophosphocholine esterase) (LDL-associated phospholipase A2) (LDL-PLA(2)) (PAF 2-acylhydrolase) | Pla2g7 Pafah | Mus musculus (Mouse) | 440 |
| Q62266 | Q62266 | reviewed | SPR1A_MOUSE | Cornifin-A (Small proline-rich protein 1A) (SPR1 A) (SPR1A) | Sprr1a | Mus musculus (Mouse) | 144 |
| E9Q557 | E9Q557 | reviewed | DESP_MOUSE | Desmoplakin (DP) | Dsp | Mus musculus (Mouse) | 2883 |
| P10639 | P10639 | reviewed | THIO_MOUSE | Thioredoxin (Trx) (ATL-derived factor) (ADF) | Txn Txn1 | Mus musculus (Mouse) | 105 |
| P17742 | P17742 | reviewed | PPIA_MOUSE | Peptidyl-prolyl cis-trans isomerase A (PPIase A) (EC 5.2.1.8) (Cyclophilin A) (Cyclosporin A-binding protein) (Rotamase A) (SP18) [Cleaved into: Peptidyl-prolyl cis-trans isomerase A, N-terminally processed] | Ppia | Mus musculus (Mouse) | 164 |
| Q02257 | Q02257 | reviewed | PLAK_MOUSE | Junction plakoglobin (Desmoplakin III) (Desmoplakin-3) | Jup | Mus musculus (Mouse) | 745 |

**M2 BMDM-EV signature proteins**

| From | Entry | Reviewed | Entry Name | Protein names | Gene Names | Organism | Length |
|---|---|---|---|---|---|---|---|
| P01756 | P01756 | reviewed | HVM12_MOUSE | Ig heavy chain V region MOPC 104E | | Mus musculus (Mouse) | 117 |
| P05064 | P05064 | reviewed | ALDOA_MOUSE | Fructose-bisphosphate aldolase A (EC 4.1.2.13) (Aldolase 1) (Muscle-type aldolase) | Aldoa Aldo1 | Mus musculus (Mouse) | 364 |
| P16110 | P16110 | reviewed | LEG3_MOUSE | Galectin-3 (Gal-3) (35 kDa lectin) (Carbohydrate-binding protein 35) (CBP 35) (Galactose-specific lectin 3) (IgE-binding protein) (L-34 galactoside-binding lectin) (Laminin-binding protein) (Lectin L-29) (Mac-2 antigen) | Lgals3 | Mus musculus (Mouse) | 264 |
| P62242 | P62242 | reviewed | RS8_MOUSE | Small ribosomal subunit protein eS8 (40S ribosomal protein S8) | Rps8 | Mus musculus (Mouse) | 208 |
| P62806 | P62806 | reviewed | H4_MOUSE | Histone H4 | H4c1 Hist1h4a; H4c2 H4-53 Hist1h4b; H4c3 H4-12 Hist1h4c; | Mus musculus (Mouse) | 103 |
| Q6GQT1 | Q6GQT1 | reviewed | A2MG_MOUSE | Alpha-2-macroglobulin-P (Alpha-2-macroglobulin) | A2m A2mp | Mus musculus (Mouse) | 1474 |
| O35744 | O35744 | reviewed | CHIL3_MOUSE | Chitinase-like protein 3 (EC 3.2.1.52) (Beta-N-acetylhexosaminidase Ym1) (Chitinase-3-like protein 3) (ECF-L) (Eosinophil chemotactic cytokine) (Secreted protein Ym1) | Chil3 Chi3l3 Ym1 | Mus musculus (Mouse) | 398 |
| O70435 | O70435 | reviewed | PSA3_MOUSE | Proteasome subunit alpha type-3 (Macropain subunit C8) (Multicatalytic endopeptidase complex subunit C8) (Proteasome component C8) (Proteasome subunit K) (Proteasome subunit alpha-7) (alpha-7) | Psma3 | Mus musculus (Mouse) | 255 |
| O88342 | O88342 | reviewed | WDR1_MOUSE | WD repeat-containing protein 1 (Actin-interacting protein 1) (AIP1) | Wdr1 | Mus musculus (Mouse) | 606 |
| P04117 | P04117 | reviewed | FABP4_MOUSE | Fatty acid-binding protein, adipocyte (3T3-L1 lipid-binding protein) (Adipocyte lipid-binding protein) (ALBP) (Adipocyte-type fatty acid-binding protein) (A-FABP) (AFABP) (Fatty acid-binding protein 4) (Myelin P2 protein homolog) (P15) (P2 adipocyte protein) (Protein 422) | Fabp4 Ap2 | Mus musculus (Mouse) | 132 |
| P09066 | P09066 | reviewed | HME2_MOUSE | Homeobox protein engrailed-2 (Homeobox protein en-2) (Mo-En-2) | En2 En-2 | Mus musculus (Mouse) | 324 |
| P10107 | P10107 | reviewed | ANXA1_MOUSE | Annexin A1 (Annexin I) (Annexin-1) (Calpactin II) (Calpactin-2) (Chromobindin-9) (Lipocortin I) (Phospholipase A2 inhibitory protein) (p35) [Cleaved into: Annexin Ac2-26] | Anxa1 Anx1 Lpc-1 Lpc1 | Mus musculus (Mouse) | 346 |
| P12265 | P12265 | reviewed | BGLR_MOUSE | Beta-glucuronidase (EC 3.2.1.31) | Gusb Gus Gus-s | Mus musculus (Mouse) | 648 |
| P13020 | P13020 | reviewed | GELS_MOUSE | Gelsolin (Actin-depolymerizing factor) (ADF) (Brevin) | Gsn Gsb | Mus musculus (Mouse) | 780 |
| P14069 | P14069 | reviewed | S10A6_MOUSE | Protein S100-A6 (5B10) (Calcyclin) (Prolactin receptor-associated protein) (S100 calcium-binding protein A6) | S100a6 Cacy | Mus musculus (Mouse) | 89 |
| P16045 | P16045 | reviewed | LEG1_MOUSE | Galectin-1 (Gal-1) (14 kDa lectin) (Beta-galactoside-binding lectin L-14-I) (Galaptin) (Lactose-binding lectin 1) (Lectin galactoside-binding soluble 1) (S-Lac lectin 1) | Lgals1 Gbp | Mus musculus (Mouse) | 135 |
| P17742 | P17742 | reviewed | PPIA_MOUSE | Peptidyl-prolyl cis-trans isomerase A (PPIase A) (EC 5.2.1.8) (Cyclophilin A) (Cyclosporin A-binding protein) (Rotamase A) (SP18) [Cleaved into: Peptidyl-prolyl cis-trans isomerase A, N-terminally processed] | Ppia | Mus musculus (Mouse) | 164 |
| P20060 | P20060 | reviewed | HEXB_MOUSE | Beta-hexosaminidase subunit beta (EC 3.2.1.52) (Beta-N-acetylhexosaminidase subunit beta) (Hexosaminidase subunit B) (N-acetyl-beta-glucosaminidase subunit beta) | Hexb | Mus musculus (Mouse) | 536 |
| P20152 | P20152 | reviewed | VIME_MOUSE | Vimentin | Vim | Mus musculus (Mouse) | 466 |
| P26041 | P26041 | reviewed | MOES_MOUSE | Moesin (Membrane-organizing extension spike protein) | Msn | Mus musculus (Mouse) | 577 |
| P40124 | P40124 | reviewed | CAP1_MOUSE | Adenylyl cyclase-associated protein 1 (CAP 1) | Cap1 Cap | Mus musculus (Mouse) | 474 |
| P40142 | P40142 | reviewed | TKT_MOUSE | Transketolase (TK) (EC 2.2.1.1) (P68) | Tkt | Mus musculus (Mouse) | 623 |
| P48036 | P48036 | reviewed | ANXA5_MOUSE | Annexin A5 (Anchorin CII) (Annexin V) (Annexin-5) (Calphobindin I) (CPB-I) (Endonexin II) (Lipocortin V) (Placental anticoagulant protein 4) (PP4) (Placental anticoagulant protein I) (PAP-I) (Thromboplastin inhibitor) (Vascular anticoagulant-alpha) (VAC-alpha) | Anxa5 Anx5 | Mus musculus (Mouse) | 319 |
| P48678 | P48678 | reviewed | LMNA_MOUSE | Prelamin-A/C [Cleaved into: Lamin-A/C] | Lmna Lmn1 | Mus musculus (Mouse) | 665 |
| P61982 | P61982 | reviewed | 1433G_MOUSE | 14-3-3 protein gamma [Cleaved into: 14-3-3 protein gamma, N-terminally processed] | Ywhag | Mus musculus (Mouse) | 247 |
| P62814 | P62814 | reviewed | VATB2_MOUSE | V-type proton ATPase subunit B, brain isoform (V-ATPase subunit B 2) (Endomembrane proton pump 58 kDa subunit) (Vacuolar proton pump subunit B 2) | Atp6v1b2 Atp6b2 Vat2 | Mus musculus (Mouse) | 511 |
| P62897 | P62897 | reviewed | CYC_MOUSE | Cytochrome c, somatic | Cycs | Mus musculus (Mouse) | 105 |
| P62962 | P62962 | reviewed | PROF1_MOUSE | Profilin-1 (Profilin I) | Pfn1 | Mus musculus (Mouse) | 140 |
| Q01853 | Q01853 | reviewed | TERA_MOUSE | Transitional endoplasmic reticulum ATPase (TER ATPase) (EC 3.6.4.6) (15S Mg(2+)-ATPase p97 subunit) (Valosin-containing protein) (VCP) | Vcp | Mus musculus (Mouse) | 806 |
| Q08857 | Q08857 | reviewed | CD36_MOUSE | Platelet glycoprotein 4 (Glycoprotein IIIb) (GPIIIB) (PAS IV) (PAS-4) (Platelet glycoprotein IV) (GPIV) (CD antigen CD36) | Cd36 | Mus musculus (Mouse) | 472 |
| Q60692 | Q60692 | reviewed | PSB6_MOUSE | Proteasome subunit beta type-6 (EC 3.4.25.1) (Low molecular mass protein 19) (Macropain delta chain) (Multicatalytic endopeptidase complex delta chain) (Proteasome delta chain) (Proteasome subunit Y) (Proteasome subunit beta-1) (beta-1) | Psmb6 Lmp19 | Mus musculus (Mouse) | 238 |
| Q61171 | Q61171 | reviewed | PRDX2_MOUSE | Peroxiredoxin-2 (EC 1.11.1.24) (Thiol-specific antioxidant protein) (TSA) (Thioredoxin peroxidase 1) (Thioredoxin-dependent peroxide reductase 1) (Thioredoxin-dependent peroxiredoxin 2) | Prdx2 Tdpx1 Tpx | Mus musculus (Mouse) | 198 |
| E9PVG8 | E9PVG8 | unreviewed | E9PVG8_MOUSE | Mucin-2 | Fcgbpl1 9530053A07Rik | Mus musculus (Mouse) | 2581 |
| O70370 | O70370 | reviewed | CATS_MOUSE | Cathepsin S (EC 3.4.22.27) | Ctss Cats | Mus musculus (Mouse) | 340 |
| P01837 | P01837 | reviewed | IGKC_MOUSE | Immunoglobulin kappa constant (Ig kappa chain C region MOPC 21) | Igkc | Mus musculus (Mouse) | 107 |
| P01942 | P01942 | reviewed | HBA_MOUSE | Hemoglobin subunit alpha (Alpha-globin) (Hemoglobin alpha chain) [Cleaved into: Hemopressin] | Hba Hba-a1 | Mus musculus (Mouse) | 142 |
| P02088 | P02088 | reviewed | HBB1_MOUSE | Hemoglobin subunit beta-1 (Beta-1-globin) (Hemoglobin beta-1 chain) (Hemoglobin beta-major chain) | Hbb-b1 | Mus musculus (Mouse) | 147 |
| P07724 | P07724 | reviewed | ALBU_MOUSE | Albumin | Alb Alb-1 Alb1 | Mus musculus (Mouse) | 608 |
| P08905 | P08905 | reviewed | LYZ2_MOUSE | Lysozyme C-2 (EC 3.2.1.17) (1,4-beta-N-acetylmuramidase C) (Lysozyme C type M) | Lyz2 Lyz Lyzs | Mus musculus (Mouse) | 148 |
| P11680 | P11680 | reviewed | PROP_MOUSE | Properdin (Complement factor P) | Cfp Pfc | Mus musculus (Mouse) | 464 |
| P21460 | P21460 | reviewed | CYTC_MOUSE | Cystatin-C (Cystatin-3) | Cst3 | Mus musculus (Mouse) | 140 |
| P29391 | P29391 | reviewed | FRIL1_MOUSE | Ferritin light chain 1 (Ferritin L subunit 1) | Ftl1 Ftl Ftl-1 | Mus musculus (Mouse) | 183 |

**Supplemental Table 1.** Signature EV protein identities for M1 and M2 BMDMs. The workflow is reflected by Supplemental Figure 3.
